# Oxygen-generated spatial distribution of cell population links to p53 status

**DOI:** 10.1101/2019.12.27.889394

**Authors:** Shashank Taxak, Uttam Pati

## Abstract

Low oxygen induces wild type p53 inactivation and selects for mutant-like p53 phenotypes for aggressive tumor growth. Recently, we have shown wild type p53 as a cellular oxygen-sensor that operates in switch-like fashion to transform its characters of a tumor suppressor or promoter in a gradient of hypoxia. However, it is unclear how hypoxic tumors select for wild type p53 phenotypes for oxygen-sensitive responses. Here, we show that oxygen-generated spatial distribution of the cell population induces p53 phenotype-specific survival or death. We have found that a dynamic state of spatial scatters or clustering patterns of cell populations favor the survival of wild type more than the mutant phenotypes in a wide range of oxygen fluctuation by affecting p53 subcellular localization. Our results demonstrate how spatial distribution could function to establish wild type p53-mediated oxygen sensing and cell fate decisions in a cell population with heterogeneous p53 allele status. We anticipate that such behavior of cells in a gradient of oxygen can be utilized by the hypoxic tumors to maintain distinct p53 alleles and determine the release and metastasis of single or clustered circulating tumor cells (CTCs).

**Summary sentence:** Oxygen variation results in p53 phenotype-specific cell fate via the spatial distribution pattern of the cell population

## INTRODUCTION

The p53 status is quite heterogeneous in cancer (1). Tumors usually acquire a mutant (MT) status of the p53 gene which introduces either loss- (LOF) or gain-of-function (GOF). Eventually, the two effects overcome p53’s tumor suppressive and introduce the tumor-promoting activity and make cancer more aggressive. In addition to the dominant-negative effect, the MT allele might also induce loss of heterozygosity (LOH) in which the wild type (WT) allele at a given locus is either deleted or mutated (2). Despite these effects, a large fraction of cancer types also carries on p53WT or null phenotypes. Restoration of WT function has led to the regression of different MT p53 tumor types in mice (3, 4). Conversely, WT p53 may also acquire the gain of pro-survival function in cancer cells that counteract tumor therapy (5). In contradiction to LOF or GOF, MT p53 might gain an alternate pathway for apoptosis (6). Moreover, cells may amplify phosphatidylinositol5-phosphate4-kinase type-2α (PIP4K2α) and PIP4K2β genes to survive in the absence of p53 (7). These findings suggest that the functions of p53 are strictly dependent upon its allele’s status and tumors must continuously evolve mechanisms to select for more appropriate phenotypes for unrestricted growth. Oxygen is a major limiting factor for cell survival. Without a proper vascular network of their own, tumors cannot grow beyond certain limits (8). It is known that a gradient of low oxygen selects for p53 mutant-like phenotypes and affects p53’s status of a tumor suppressor by diverse mechanisms (9–13). Recently, we have shown WT p53 as a cellular oxygen sensor that operates through the tetramer→octamer switch in its homo-oligomerization system and controls survival or death in a gradient of hypoxia (14). We asked how the same gradient selects or maintains WT p53 phenotypes in a heterogeneous cancer cell population.

## RESULTS

### Variable oxygen generates distinct spatial distribution patterns of the cell population

A two-dimensional layer of proliferating HCT116 ^p53WT/WT^, MDA-MB231 ^p53MT/MT^, HCT116 ^p53-/-^ or DU145 ^p53WT/MT^ cell line suggested scattered (SP) and clustered patterns (CP) of the spatial distribution of a cell population (Fig. 1A). Followed by gentle detachment from culture dishes to minimize cluster disintegration, flow cytometry measured forward (FSC) and side scatter (SSC) which distinguished single (low FSC) and aggregated cells (high FSC) based on size and complexity. The measurements differentiated SP and CP in a population. The FSC v/s SSC scatter plots showed the oxygen-dependent dynamics of SP⇄CP transitions. In normoxia, almost an equal fraction of HCT116 ^p53WT/WT^, DU145 ^p53WT/MT^, MDA-MB231 ^p53MT/MT^ resided in both SP and CP. However, an enhanced fraction of HCT116 ^p53-/-^cells followed CP (Fig 1B, *left panel*). This indicates that p53 might be responsible for an equilibrium state of SP⇄CP for the stability of both the patterns under normal growth conditions irrespective of its allele’s status in the cell. After transfer to hypoxia, HCT116 ^p53WT/WT^ cells showed SP→CP shift. However, MDA-MB231 ^p53MT/MT^ or HCT116 ^p53-/-^ cells underwent a reverse SP⇄CP shift. Interestingly, DU145 ^p53WT/MT^ maintained SP⇄CP state (Fig 1B, *middle panel*). Upon reoxygenation, the observed shifts in hypoxia were reversed for each cell line (Fig 1B, *right panel*). Although oxygen can induce shifts in SP⇄CP state without p53, as observed for HCT116^p53-/-^ cells in normoxia, hypoxia or reoxygenation condition, a closer examination reveals that it variably modulates SP⇄CP for WT and MT p53 phenotypes depending upon the concentration.

**Figure 1.**
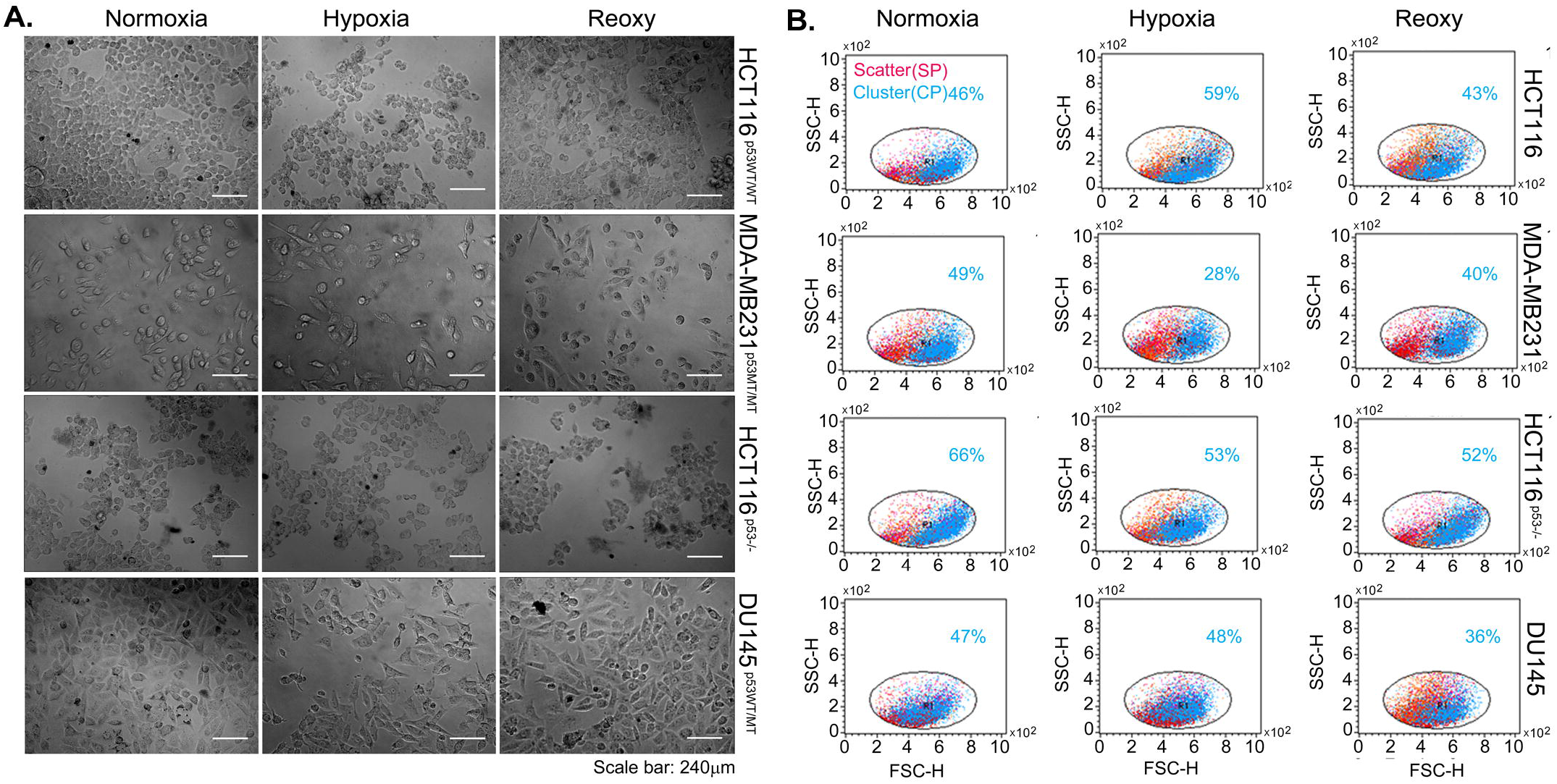
Oxygen-dependent spatial distribution pattern of the cell population. **A.** Light microscope images show SP and CP in a cell population under normoxia (5% O_2_), hypoxia (1% O_2_) or reoxygenation (21% O_2_). **B.** Flow cytometry-based detection of dynamics of the spatial distribution of cells. 10,000 cells were analyzed for SSC v/s FSC plot generation by flow cytometry.

### Spatial distribution patterns favor p53 wild type phenotypes

Next, we checked whether such oxygen-generated spatial distribution patterns select the p53 phenotypes via survival or death. For this purpose, first, we analyzed cell fate associated with SP or CP in different oxygen conditions. For this task, we adapted a flow cytometry assay based on the principle described elsewhere (15). Together Propidium iodide and YO-PRO1 (termed as PY) stains both the nucleus and cytoplasm of the cells (see Material and methods). PY-stained cells differ in both pulse width (W) and area (A) when present in SP and CP. Upon excitation by flow cytometer, the fraction of cells with PI fluorescence of Low W & high A represents the extent of death in SP and inversely, high W & low A fractions correspond to death in CP (Fig. S1). Trypan blue staining of the monolayer cultures provides a visual authentication of our adaptation of the technique by light microscopy. Cell fate associated with SP or CP showed that SP allows selection of p53 positive by eliminating p53 null phenotypes by death in normal growth conditions. By contrast, CP selects for p53 positive and eliminates p53 null phenotypes in hypoxia. Interestingly, SP and CP protect WT and MT p53 along with p53 null phenotypes respectively upon reoxygenation (Fig. 2). Immunofluoro-cytostaining shows that cytoplasmic p53 is favored for the selection by CP in hypoxia. However, nuclear and cytoplasmic p53 are preferred for the selection of WT and MT phenotypes by SP and CP upon reoxygenation respectively (Fig S2). Next, to decipher the role of oxygen-generated spatial distribution patterns in the selection of WT and MT phenotypes, we compared FSC v/s SSC and PY-W v/s PY-A plots. Although CP selects p53 positive cells, hypoxia generates SP→CP for WT and SP←CP for MT and null phenotype selection. Similarly, reoxygenation stimulates SP←CP for WT and SP→CP for MT phenotypes. The shifts in SP⇄CP try to maintain p53 null and p53WT/MT phenotypes irrespective of oxygen variation (Fig 3A). Strikingly, the shifts specifically favored the survival of WTp53 phenotypes in a wide range of oxygen fluctuation. This indicates oxygen-generated spatial distribution patterns as a mechanism to select for WT species for oxygen-sensitive responses.

**Figure2.**
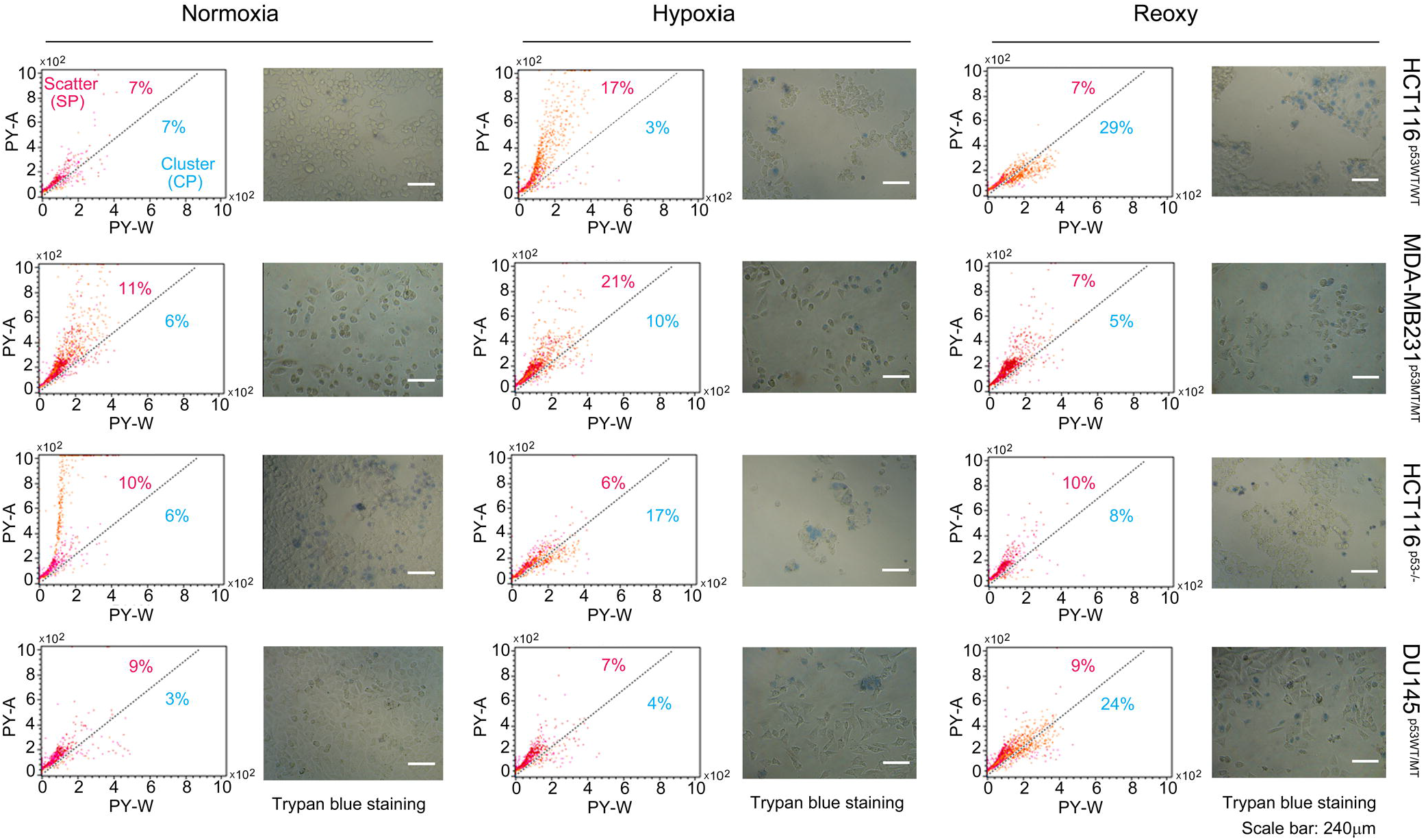
Spatial distribution affects cell fate. PY-W v/s PY-A plots show cell death or survival in SP or CP. Trypan blue staining authenticates flow cytometry-based determination of cell death in SP or CP. 10,000 cells were analyzed for PY-W v/s PY-A plot generation by Flow cytometry.

**Figure3.**
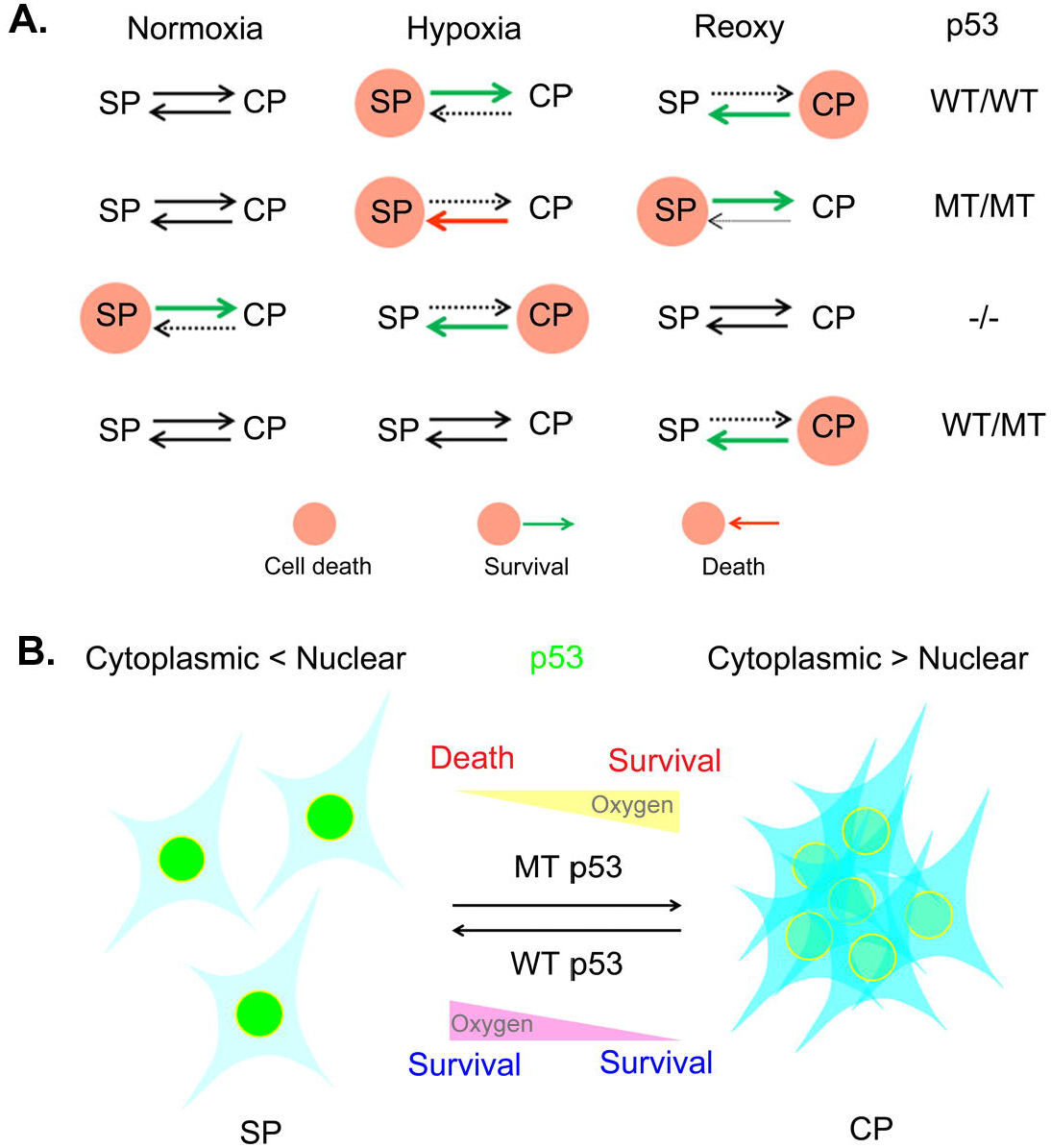
The oxygen-generated spatial distribution of cells induces p53 phenotype-specific cell fate. **A.** Selection of different p53 phenotypes by survival or death of the cells via distinct spatial distribution patterns. **B.** The model of the study. Oxygen gradient represents hypoxia (low) and reoxygenation or normoxia (high) conditions.

## DISCUSSION

Here, we report our discovery that oxygen controls the spatial distribution pattern of the cell population to decide the survival or death of the distinct p53 phenotypes. By using human cancerous cell lines with p53WT/WT, MT/MT, -/- and WT/MT allele’s status and Flow cytometry-based assays at different oxygen levels, we show that variation in oxygen levels triggers either scatter or clustering patterns of the cells depending upon the p53 status. Such dynamic arrangements transform the death or survival potential of WT and MT p53 phenotypes. p53 subcellular localization is linked to cell survival or death (13, 16–19) and differ with SP and CP. Under fluctuating oxygen, SP and CP tend to maintain diminished and enhanced cytoplasmic to nuclear p53 distribution ratio respectively which might result in mislocalization of WT or MT p53 and transformation of cell fate upon SP⇄CP transitions. Enhanced survival of WT p53 cells in a wide range of oxygen variation justifies the significance of such spatial dynamics in selecting WT p53 for oxygen-sensitive responses (14) (Fig 3B).

An additional implication of such a mechanism can be anticipated in the metastasis of cells. It is known that solid tumors can release a high number of circulating tumor cells (CTC) into the circulation. CTC could be a single or cluster of cells that initiate metastasis. A clustered CTC is more metastatic than a single CTC (20–22). It is evident that the mutant p53 phenotype is selected mostly by CP which might explain the enhanced metastatic tendency of clustered CTC (23). Our findings link oxygen and spatial clustering of cells that might control the release of single or clustered CTCs and determine their metastasis based on the p53 allele’s status.

In conclusion, our study reveals a unique model of biological oxygen sensing in which cells respond to fluctuating oxygen by selecting WT p53 phenotypes in a cell population to transform cell fate. Further investigations might reveal how this model of oxygen sensing influences the molecular dynamics of intracellular oxygen sensors p53 or HIF-1α to drive cancer progression in a gradient of hypoxia.

## EXPERIMENTAL PROCEDURES

### Cell culture and Hypoxia treatment

HCT116 ^p53+/+^, HCT116 ^p53-/-^, MDA-MB231 ^p53MT/MT^ (NCCS, Pune), A431 ^p53MT/MT^(NCCS, Pune), and DU145 p53^WT/MT^ (NCCS, Pune) were cultured in Dulbecco’s modified Eagle’s medium (Merck, D5648) supplemented with 10% (v/v) FCS (Thermofisher, 10438018), penicillin (100 U/ml), streptomycin (100 U/ ml) and L-glutamine (4 mM) (Merck, G3126), pH 7.4. For the proliferating cell cultures, 5% CO_2_ was maintained in a humidified incubator at 37 °C. Hypoxia exposure to the cell culture was performed under a humidified condition in a hypoxia chamber (Plas Labs inc., USA) equipped with O_2_ and CO_2_ sensors. O_2_ and CO_2_ levels were quickly restored upon the detection of fluctuations (by ±0.2 units) from the set levels through the automated purging of gases by inbuilt sensors. The normal atmospheric pressure was maintained through the N_2_ gas. Similarly, the integrated thermostat maintained the temperature at 37°C. Just before the exposure, DMEM was replenished in the cultures. Re-oxygenation was performed by transferring the cultures from the hypoxia chamber to the CO2 (5%) incubator for a 24h period.

### Flow cytometry

Following the treatments, cells were rinsed with PBS and harvested by mild trypsinization for 10-15 seconds at 37°C. The trypsinized cells were gently detached with chilled PBS supplemented with 2% FBS. The cells were gently washed and diluted (approx. 30,000 cells per ml) in PBS with 2% FBS to minimize cellular clumping. Vybrant apoptosis assay kit (Thermofisher, V13243) was used to stain cells simultaneously with PI and YO-PRO1 as per manufacturer’s instruction. PI-H v/s YO-PRO1-H scatter plot was used for gating single and clustered cells in FSC v/s SSC plot or dead fraction of SP and CP in PI-W v/s PI-A plot using BD CellQuest Pro software. The cells were vortexed gently prior to the flow cytometry on a BD FACS Calibur (BD Biosciences, USA). 488nm blue-laser was used to excite both YO-PRO1 and the PI and the signals were captured in FL1 for YO-PRO1 and FL2 channels for PI emissions.

### Gating strategy

FL2 channel captures PI maximum emission and YO-PRO1 bleed-through due to which their simultaneous emissions in FL2 are termed as PY. The maximum emission of YO-PRO1 is captured in the FL1 channel. YO-PRO1 can distribute in both cytoplasm and nucleus of apoptotic cells. In clusters, a single apoptotic cell may allow distribution of YO-PRO1 in the cytoplasm of other aggregated cells through cytoplasmic connections but PI will dominantly stain the nucleus of single cells disintegrated from clusters. Thus, YO-PRO1 (FL1-H) v/s PI (FL2-H) scatter plot can be used to gate and differentiate single and clustered cells in the FSC v/s SSC plot. Also, PY-W v/s PY-A would differentiate death in SP or CP. We validated our findings through visualization by light microscopy and trypan blue staining.

### Trypan blue staining

After treatments, cells were rinsed with PBS. A 50% trypan blue solution was diluted in PBS was applied to the monolayer of cells for 5 min. Cells were washed once and fresh PBS was added in the culture dishes before imaging by light microscope (Nikon, TMS-F, Japan).

### Immunocytostaining

The experimental procedure was performed as described previously (14). Cells were grown on coverslips and exposed to different conditions of oxygen. Exposed cells were washed in PBS and chemically fixed with 4% (v/v) paraformaldehyde at room temperature (10 min). 1mM glycine was added to quench the reaction and cells were again washed thrice in PBS. Cells were then treated with 100% chilled methanol for permeabilization which followed washing in PBS to completely remove methanol. Autofluorescence was removed by sodium borohydride treatment for 20 min at room temperature. Again cells were washed and blocked in 4% BSA in PBS (with 0.1% TX-100) overnight at 4°C. Endogenous p53 was detected by anti-p53DO1 primary antibody (1:1000 dilutions) at 22°C for 4h. Indirect immune labeling was performed by FITC-tagged secondary antibody (1:5000 dilutions) for 2h in dark at room temperature. From now onwards all procedures were performed in dark. Secondary antibodies were washed and SlowFade antifade kit (Thermofisher, S2828) was used to mount the coverslips for Confocal imaging by Zeiss LSM510 (Germany) microscope equipped with 60X oil immersion objective (NA 1.4). FITC was excited by 488nm laser and emission was captured by 530-600nm BP filter.

## Acknowledgments

S Das, HCT116 ^p53+/+^ and HCT116 ^p53-/-^ cells; P Malhotra for support in Flow cytometry.

## Author contributions

ST conceived the idea; ST conducted experiments; ST and UP analyzed data; ST wrote the manuscript.

## Data and materials availability

All data is available in the main text or the supplementary figures.

## FOOTNOTES

Funding was provided by the Indian council of medical research (ICMR), Govt. of India through UPOE grants, as well as Ph.D. Fellowship to S.T. from Council of Scientific and Industrial Research (CSIR), Govt.of India.

## Abbreviations

p53: 53kDa tumor protein
SP: Scattered pattern
CP: Clustered pattern
WT: Wild type
MT: Mutant
FSC: forward scatter
SSC: side scatter
CTC: circulating tumor cells

