## Supplementary figures for "Oxygen-generated spatial distribution of cell population links to p53 status"

Shashank Taxak, *Uttam Pati
Transcriptional & Human Biology Laboratory, School of Biotechnology, Jawaharlal Nehru University, New Delhi-110067, India

Authors' address



### This pdf file includes:

Fig S1, S2

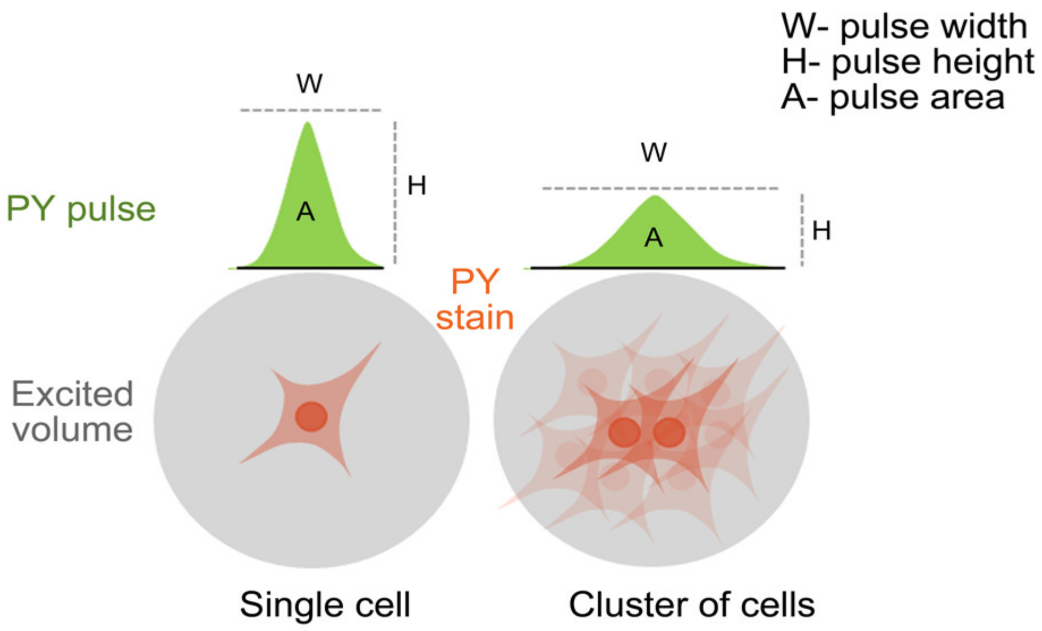


**Figure S1. Principle of flow cytometry-based detection of cell death in SP and CP.** Death in single and aggregated cells is detected by low and high PY-W fractions respectively. PY-H or PY-A may vary depending upon PY internalization by the cells or number of cells undergoing death individually or in a cluster.

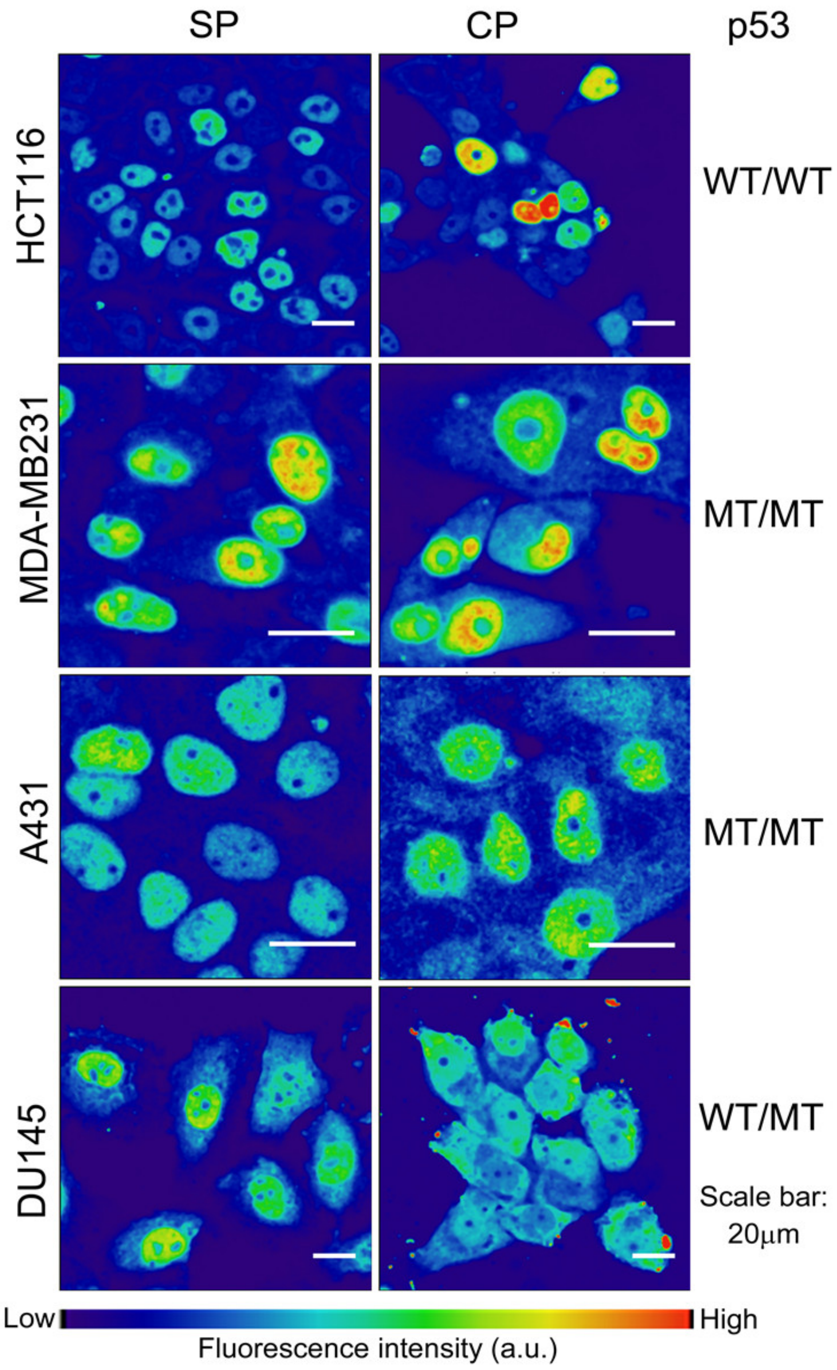


**Figure S2. The spatial distribution pattern of the cell population affects the subcellular localization of p53.** Fluorescent images are pseudocolored and color calibration bar indicates fluorescence intensity (a.u.)
